# Compression and *k*-mer based Approach For Anticancer Peptide Analysis

**DOI:** 10.1101/2024.10.05.616787

**Authors:** Sarwan Ali, Tamkanat E Ali, Prakash Chourasia, Murray Patterson

## Abstract

Our research delves into the imperative realm of anti-cancer peptide sequence analysis, an essential domain for biological researchers. Presently, neural network-based methodologies, while exhibiting precision, encounter challenges with a substantial parameter count and extensive data requirements. The recently proposed method to compute the pairwise distance between the sequences using the compression-based approach [26] focuses on compressing entire sequences, potentially overlooking intricate neighboring information for individual characters (i.e., amino acids in the case of protein and nucleotide in the case of nucleotide) within a sequence. The importance of neighboring information lies in its ability to provide context and enhance understanding at a finer level within the sequences being analyzed. Our study advocates an innovative paradigm, where we integrate classical compression algorithms, such as Gzip, with a pioneering *k*-mers-based strategy in an incremental fashion. Diverging from conventional techniques, our method entails compressing individual *k*-mers and incrementally constructing the compression for subsequences, ensuring more careful consideration of neighboring information for each character. Our proposed method improves classification performance without necessitating custom features or pre-trained models. Our approach unifies compression, Normalized Compression Distance, and *k*-mers-based techniques to generate embeddings, which are then used for classification. This synergy facilitates a nuanced understanding of cancer sequences, surpassing state-of-the-art methods in predictive accuracy on the Anti-Cancer Peptides dataset. Moreover, our methodology provides a practical and efficient alternative to computationally demanding Deep Neural Networks (DNNs), proving effective even in low-resource environments.

## 1 Introduction

Cancer is one of the leading contributors to global mortality trends [36]. Early and accurate detection of cancer can lead to timely treatment and, in turn, can save precious human lives. Sequence analyses help improve our understanding of cancer biology, including tumorigenesis, metastasis, and drug resistance mechanisms, driving further research and innovation in cancer prevention, diagnosis, and treatment [32]. The development of advanced computational techniques leads to the effective use of Anticancer peptides (ACPs) in the treatment of cancer. ACPs belong to the antimicrobial peptide (AMP) group that exhibits anticancer activity [14]. Analyzing ACPs properties identifies potent candidates for new treatments. Insights into ACP mechanisms of action, including cell interaction and immune modulation, are crucial for optimizing efficacy and minimizing side effects [10]. By analyzing ACPs and understanding their pharmacokinetics (how the body processes them), pharmacodynamics (how they exert their effects), and tissue distribution, researchers can optimize treatment strategies such as dosing regimens and combination therapies [14]. The analysis of anti-cancer peptides is crucial for advancing our understanding of their therapeutic potential, optimizing their efficacy and safety profiles, and ultimately developing effective cancer treatments.

Performing underlying ML tasks requires converting variable-length peptide sequences to numerical vectors using a sequence encoding technique, such as Amino Acid Composition (AAC) [1] involving frequency vector generation and/or di-peptide AAC (DAAC) based on the frequency of peptide pairs, etc. As a solution, CKSAAP [13] was proposed, which concatenates the DAAC feature vectors of *K*spaced amino acid pairs and has been successfully used in Anti Cancer peptide classification tasks [19]. The k-spaced amino acid group pairs (CKSAAGP) [37] is also used for representing ACPs based on the frequency of amino acid group pairs separated by *k* residues. However, such methods either do not generalize on different types of data (e.g., due to sub-optimal feature selection approach in [1, 13, 37] or struggle to capture complex relationships and interactions between features due to sparse representations [19]) or could be computationally expensive to learn the optimal embedding representations for the ACPs, hence limiting the predictive performance.

String kernels are a class of kernel methods that have gained increasing popularity in biological sequence analysis. Many string kernels have been proposed such as spectrum kernel, mismatch kernel [29], sparse representation classification [19], and local alignment kernel [39]. These string kernels have shown promising results in Anti-Cancer Peptide classification for example, an ML method involving Chou’s pseudo-amino acid composition (PseAAC) and local alignment kernel [22] successfully predicted ACPs similarly Kernel Sparse Representation Classifier [19] was also used in ACP classification. Although these methods provide accurate results they could cause an overfitting problem along with scalability issues making them memory intensive.

Deep Learning, especially Natural Language Processing methods have been used in forming numeric representations of antimicrobial peptides for example in [11] the sequence information is converted into digital vectors using a combination of BiLSTM, attention-residual algorithm, and BERT Encoder. The transfer learning-based pretrained biological language models along with CNN successfully generate anticancer peptide embeddings [18]. Despite the widespread use of these Neural Networks (NNs) and language models, demand a substantial number of parameters and extended training times, and are computationally expensive. Moreover, they heavily rely on large-scale training data, often unavailable for certain biological datasets.

To address these challenges we propose a novel compression-based approach involving *k*-mer strategy and NLP-based encoding to classify anti-cancer peptides using k-mers. Traditional methods, including NNs, predominantly focus on compressing entire sequences, potentially neglecting nuanced neighboring information for individual characters within a sequence. Inspired by recent advancements in compression-based approaches [26] our innovative method integrates the classical compression algorithm Gzip to compress individual encoded *k*-mers generated by NLP-based embedding and incrementally construct the compression for subsequences, ensuring a meticulous consideration of neighboring information for each character (Amino Acid). Gzip is a lossless data compression technique that gained popularity in Computational biology due to its easy integration with biological sequence analysis tools. Some prominent Compression-based distances include Normalized Compression Distance (NCD) [7], Normalized Information Distance (NID) [31], etc. Pairwise NCD is a parameter-free, feature-free, alignment-free, similarity metric based on compression. In our proposed method, we calculate NCD for each *k*-mer pair in the sequence data, which is further used to compute the distance matrix. To generate a low-dimensional numerical representation, we convert the distance matrix into a kernel matrix. Our proposed compression-based model not only overcomes the limitations of existing methods by eliminating the computational intensity associated with deep neural networks but also demonstrates efficiency in handling low-resource biological datasets where labeled data is scarce. The contributions of our study include:

1. A novel approach for identifying cancer by analyzing and classifying Anti-Cancer Peptides (ACPs) using compression-based models.
2. Our innovative approach integrates the classical Gzip compression algorithm with a pioneering k-mers-based strategy. It involves compressing individual *k*-mers and incrementally constructing the compression for subsequences, ensuring a meticulous consideration of neighboring information of each amino acid in a sequence.
3. We develop an algorithm for Distance Matrix computation, where we take a set of sequences as input and output a non-symmetric Distance matrix using Normalized Compression Distance (NCD) and Gzip compressor.
4. The proposed compression-based model eliminates the computational intensity associated with deep neural networks.
5. Leveraging Gzip compression, our approach becomes particularly advantageous in scenarios where labeled data is scarce demonstrating efficiency in handling low-resource datasets.
6. Our method addresses limitations observed in prior work related to Anti-Cancer Peptide classification, showcasing its applicability to a broader range of classification tasks.

## 2 Related Work

Anti-cancer peptides (ACPs) can be reconstructed or modified to increase their anti-cancer activity while lowering cytotoxicity as demonstrated in [41]. Usually, this entails ACP side chain modification and main chain reconstruction [16]. However, this work delves into recent advancements in their reconstruction and modification, which is not applicable in our case. In another work, ACP MS is proposed in [44], which makes use of the monoMonoKGap technique to extract properties from anticancer peptide sequences and create digital features. Sequential features or patterns or motifs and Physicochemical properties which encompass various molecular properties are used in [24]. The g-gap dipeptide components were optimized to create the sequence-based predictor known as iACP, which is presented by the authors of [12]. A lot of research has been conducted on the use of Neural Networks (NNs) in predicting anticancer peptides, for example, DeepACP a sequence-based deep learning tool [43], a Long Short-Term Neural Network model [42] with integrated binary profile features and a *k*-mer sparse matrix of the reduced amino-acid alphabet, convolutional neural network-recurrent neural network (CNN-RNN) [43]. Although these NN-based methods exhibit accurate performance, they are computationally expensive.

Natural Language Processing methods have made a significant mark in Anti Cancer Peptide (ACPs) classification including pre-trained language models such as ProteinBERT [25]. In another work [27] Protein-based transformers such as ESM, ProtBert, BioBERT, and SciBERT, have shown promising results in identifying ACPs. In a recent research [2] a FastText-based word embedding method involving a skip-gram model has been used to represent each peptide for forming the embedding on which a deep neural network (DNN) model is employed to accurately classify the ACPs. CancerGram [9] uses n-grams and random forests for predicting ACPs. UniRep [30], a Language model-based embedding is also used as a feature representation for anticancer peptides leading to improved ACP prediction. These Language model-based methods have shown improved classification results but have high memory requirements.

## 3 Proposed Approach

Our proposed approach consists of generating embedding for the ACP sequences based on *k*-mer compression and incremental NCD computation. Given a set of Sequences (S) we process each pair of these sequences represented as *s*_1_ and *s*_2_ in every iteration ultimately covering all possible pairs. The overall flow diagram of the proposed method is shown in Figure 1. Moreover, the pseudocode of our method is given in Algorithm 1.

**Figure 1:**
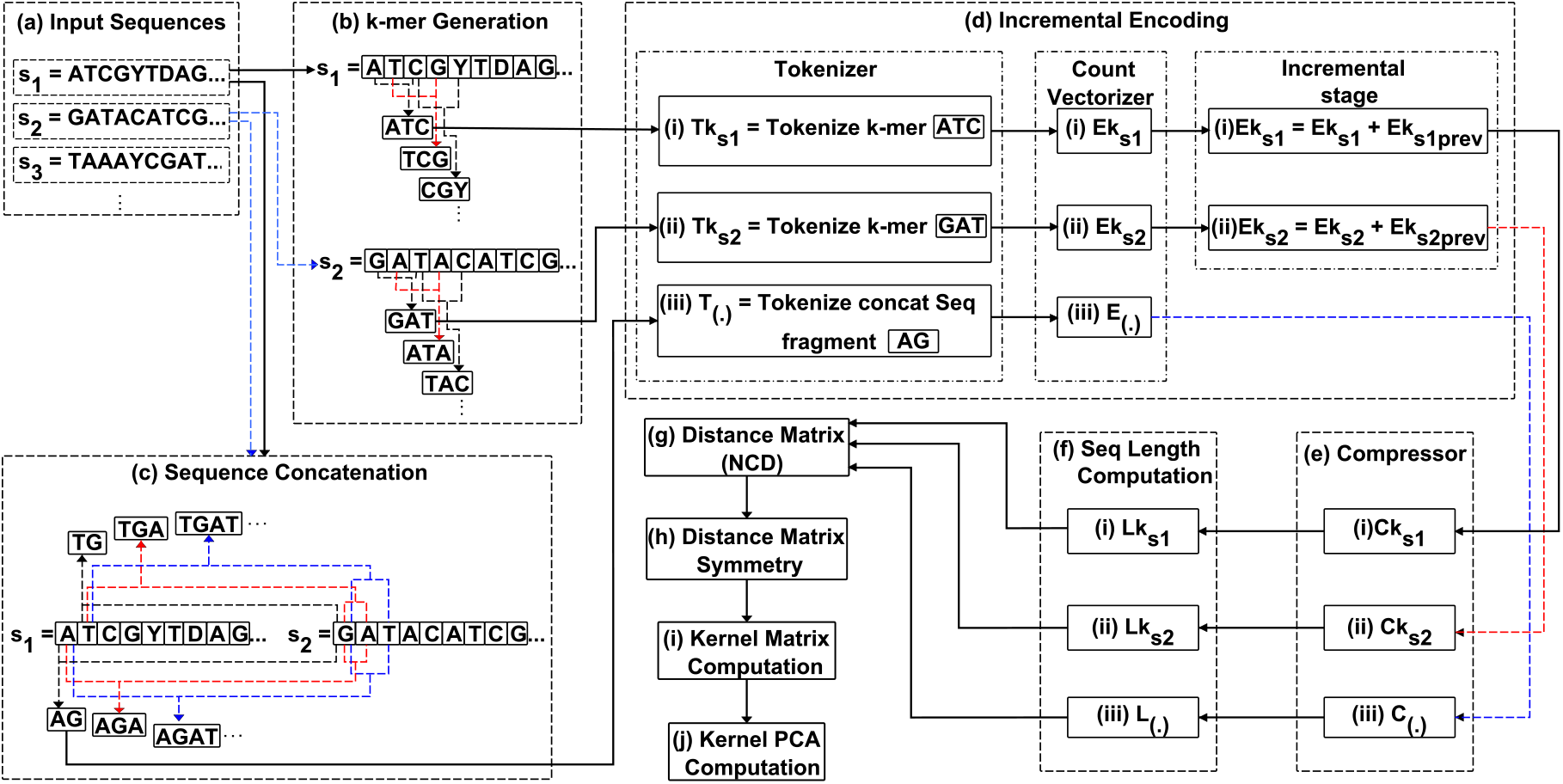
Flow diagram for the proposed approach. The figure is best seen in color.

We first compute the *k*-mers of each sequence as shown in step (b) of Figure 1 and the line numbers 10 and 15 of Algorithm 1 for *s*_1_ and *s*_2_ respectively. This is followed by encoding these *k*-mers using the function Encode in line numbers 11 and 16 which is presented in Algorithm 1 and also shown in step (d) of Figure 1, it takes in a sequence in our case *k*-mer and uses NLP based method involving tokenization of *k*-mers followed by Count Vectorization of the tokens as shown in line number 2 and 3 of Algorithm 2 to form numerical representation of the *k*-mers which is then flattened in line number 4 of Algorithm 2 and converted into a string forming the encoded version of k-mers. It can also be noticed in line numbers 11 and 16 of Algorithm 1 and incremental stage of step(d) in Figure 1 that the encoded forms of *k*-mers keep adding up in each iteration leading to incremental encoding. These incrementally encoded *k*-mers are further compressed using Gzip as shown in Algorithm 3 and step (e) of Figure 1. This is followed by calculating the lengths of these compressed incrementally encoded *k*mers in line numbers 13 and 18 for *s*_1_ and *s*_2_ respectively. To finally calculate the NCD values we need lengths of compressed concatenated sequences. In our algorithm, we adopt a unique method for concatenating by first segmenting the *s*_1_ and *s*_2_ into portions with increased length after every iteration and then concatenate in line number 19 of Algorithm 1, a detailed view of this technique is shown in step (c) of Figure 1 where the black lines show the formation of first concatenated fragments for each successive amino acid followed by red and blue lines that clearly show the increasing length of the fragments. These concatenated sequence fragments, also referred to as sub-sequences are further encoded as shown in step (d) of Figure 1 but this time without incremental stage as can be seen in line number 20 of Algorithm 1. Followed by compression and length computation stages in step(e) and step(f) of Figure 1. We then calculate NCD values using Equation 3.1.

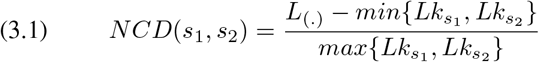

where 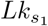 and 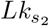 represent the lengths of the compressed incrementally encoded *k*-mers of sequences *s*_1_ and *s*_2_ respectively, and *L*_(.)_ denotes the length of the compressed form of the concatenated fragments of sequence *s*_1_ and *s*_2_. Finally, the NCD values are aggregated using the mean function to obtain a scalar value. These Scalar NCD values are appended in Distance Matrix (D) in line number 26 of Algorithm 1. This process is repeated for all pairs of sequences, resulting in the incremental encoding distance matrix, which contains the pairwise similarity values between sequences.

### Algorithm 1 Incremental Encoding Distance Matrix

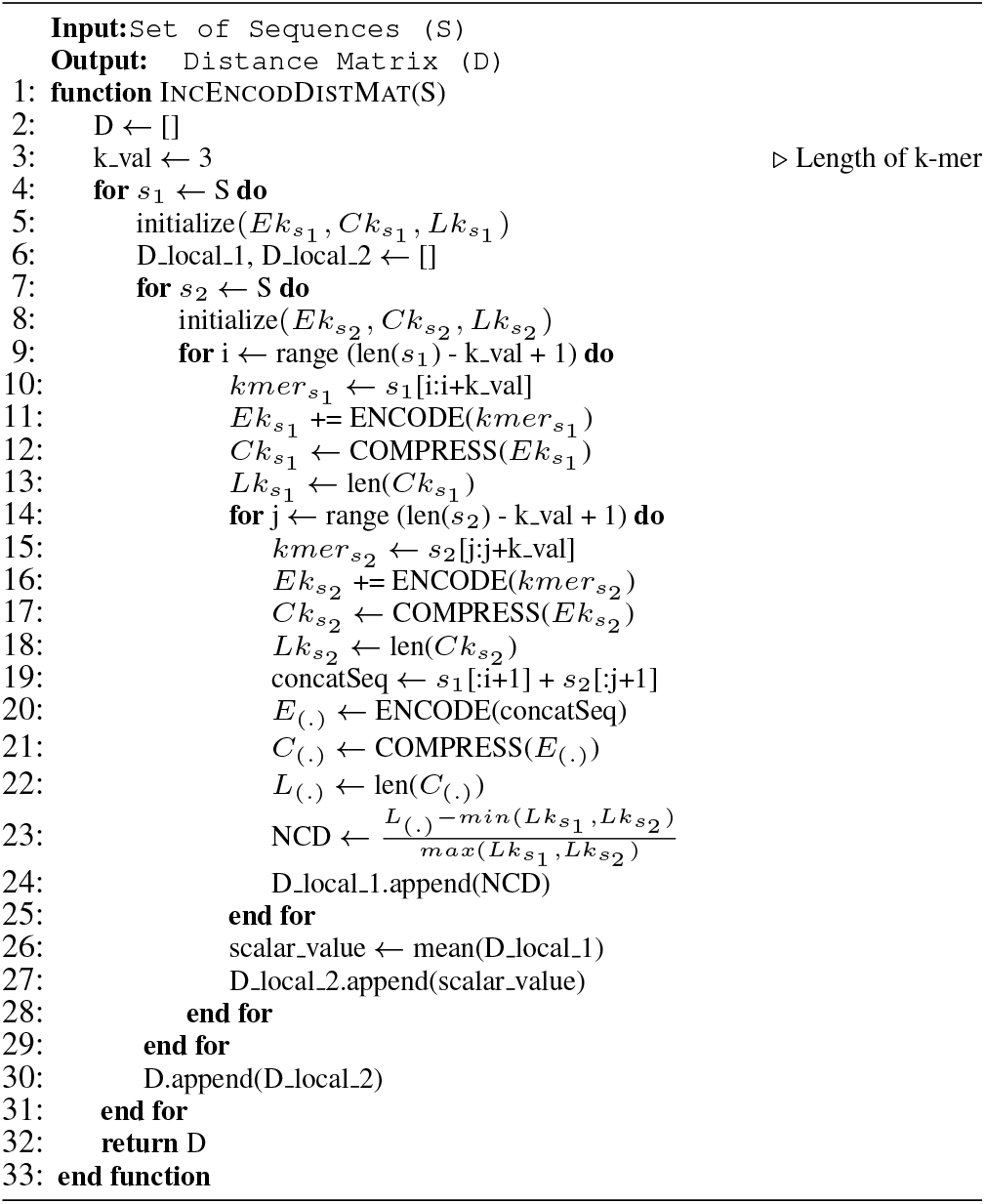

### Algorithm 2 Encoding

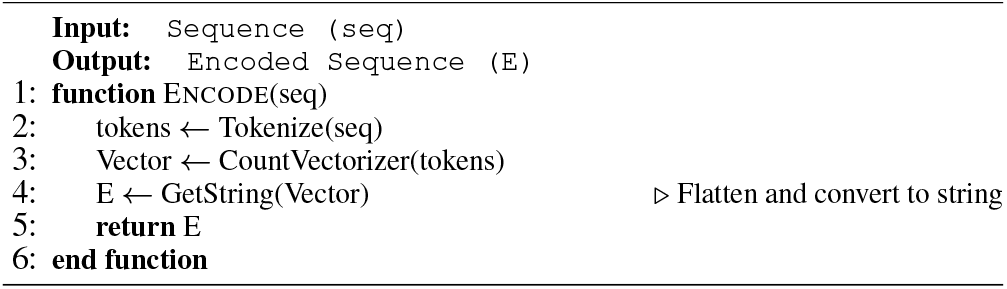

### Algorithm 3 Gzip compression

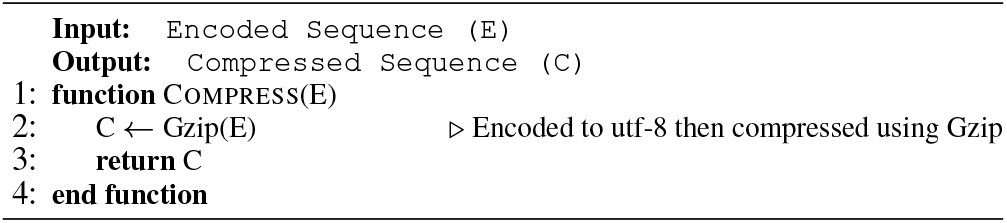

### REMARK 1.

*NCD Lower Bound: NCD values are bounded by 0 when two sequences are identical. In this case, the compressed concatenation of the sequences will be the same as the compressed length of each sequence, i*.*e. NCD*(*s*_1_, *s*_2_) = 0 *if s*_1_ = *s*_2_.

### REMARK 2.

*NCD Upper Bound: The NCD value is upper-bounded by 1 in the case where two sequences are completely dissimilar, as the compressed concatenated sequence will not provide any significant compression advantage, i*.*e. NCD*(*s*_1_, *s*_2_) *≤* 1.

### REMARK 3.

*Compression Efficiency Guarantees: The proposed algorithm’s performance depends on the efficiency of the compression algorithm (i*.*e. Gzip). Some key properties to note are the following:*

1. *Optimality in Compression: The Gzip compression algorithm ensures that common substrings or patterns within a sequence or concatenation are compressed efficiently, leading to shorter compressed lengths*.
2. *Monotonicity of Compression: The compression length for the concatenation of two sequences s*_1_ + *s*_2_ *will never exceed the sum of their individual compressed lengths, i*.*e. L*(*s*_1_ + *s*_2_) *≤ L*(*s*_1_) + *L*(*s*_2_).

*This ensures that the NCD is always well-defined*.

### REMARK 4.

*Incremental Encoding Stability: The incremental encoding approach adds k-mers to the existing sequence encoding. To ensure the stability of the algorithm, we can say the following:*

1. *Monotonic Growth in Encoding: The length of the encoded sequence grows monotonically as k-mers are added, which means the size of the encoded sequence after each iteration will never decrease. This guarantees that the compression length also grows or remains constant*.
2. *Asymptotic Compression Convergence: As the k-mer sequences are incrementally encoded and compressed, the compression lengths for large enough sequences should converge to a stable value, reflecting the overall sequence similarity*.

### 3.1 Distance Matrix Symmetry

The Distance matrix (D) obtained is of size *n × n*, where *n* represents the number of sequences in set *S*. Note that D is non-symmetric, so we convert it into a symmetric matrix by taking the average of upper and lower triangle values and replacing the original values of the matrix with the average values.

### 3.2 Kernel Matrix Computation

Then we generate a Kernel matrix from the symmetric distance matrix using a Gaussian Kernel. First, we calculate Euclidean Distance (*W*) between two pairs of distances using the equation:

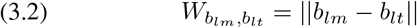

where *b*_*lm*_ and *b*_*lt*_ represent any two values of the symmetric Distance matrix.

The Gaussian Kernel (G) is defined as a measure of similarity between *b*_*lm*_ and *b*_*lt*_:

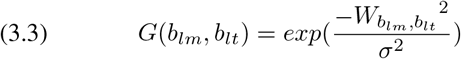

where *σ*^2^ represents the bandwidth of the kernel.

The kernel value is computed for each pair of distances in the symmetric distance matrix to get *n × n* dimensional kernel matrix using the following theory:

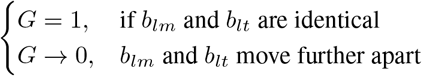

After computing the kernel matrix, we apply kernel Principal Component Analysis (PCA) to get a lower-dimensional embedding of the data. It preserves the essential information while retaining the relationships among the anti-cancer peptide sequences, including non-linear relations. This representation proves valuable for various tasks, including classification.

## 4 Experimental Setup

This section outlines the experimental setup, including details on the dataset, visualization techniques, baseline models, and metrics used for evaluation. All experiments were conducted on a computing system equipped with an Intel(R) Xeon(R) CPU E7-4850 v4 running at a clock speed of 2.10GHz and operating on a 64-bit Ubuntu OS (version 16.04.7 LTS Xenial Xerus), with a total memory capacity of 3023 GB. The algorithms were implemented using Python. The dataset was partitioned into training and testing sets using a 70 *−* 30% split ratio. To account for variability, experiments were conducted with 5 different random initializations for the train-test splits, and the results reported include both average values and standard deviations. For hyperparameter tuning, a 5-fold cross-validation strategy was employed.

### 4.1 Dataset Statistics

The dataset on Membranolytic anticancer peptides (ACPs) [21] provides details regarding peptide sequences and their corresponding anticancer effectiveness against breast and lung cancer cell lines. The target labels are classified into four groups: “very active,” “moderately active,” “experimental inactive,” and “virtual inactive.” In total, the dataset comprises 949 and 901 peptide sequences for breast and lung cancer, respectively. Table 1 shows the distribution.

**Table 1:**
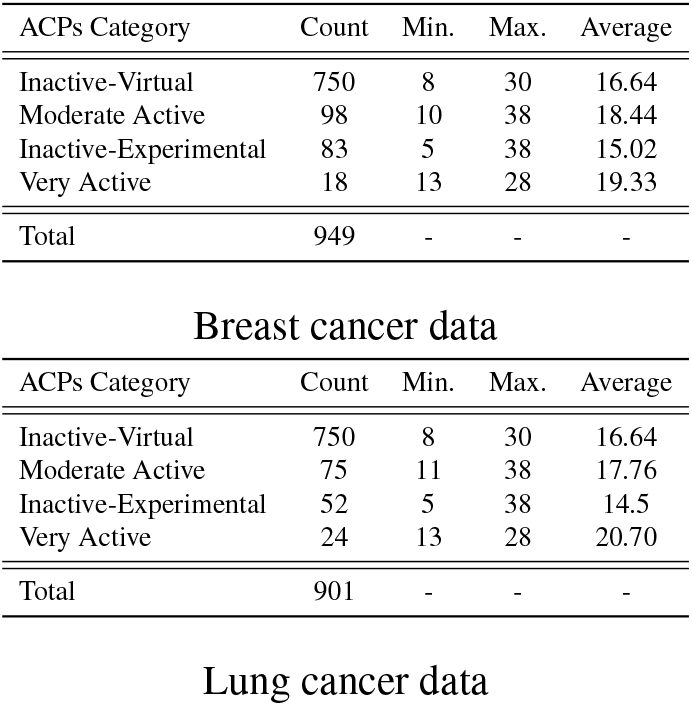
Dataset statistics for the Breast cancer and Lung cancer data. Columns represent the min., max., and average lengths of the peptide sequence.

### 4.2 Data Visualization

A widely used visualization technique, namely t-stochastic distributed neighborhood embedding (t-SNE) [15, 38], is employed to visualize the feature vectors from different embedding methods. The t-SNE plots are presented in Figure 2 and Figure 3 (in the appendix) for Breast Cancer and Lung Cancer, respectively. These plots showcase the distribution and grouping patterns of the embeddings and facilitate the qualitative assessment of how well the embeddings capture the inherent structure and relationships within the data. We can observe in Figure 2 that the proposed *k*-mers compression-based method can group similar classes reasonably well.

**Figure 2:**
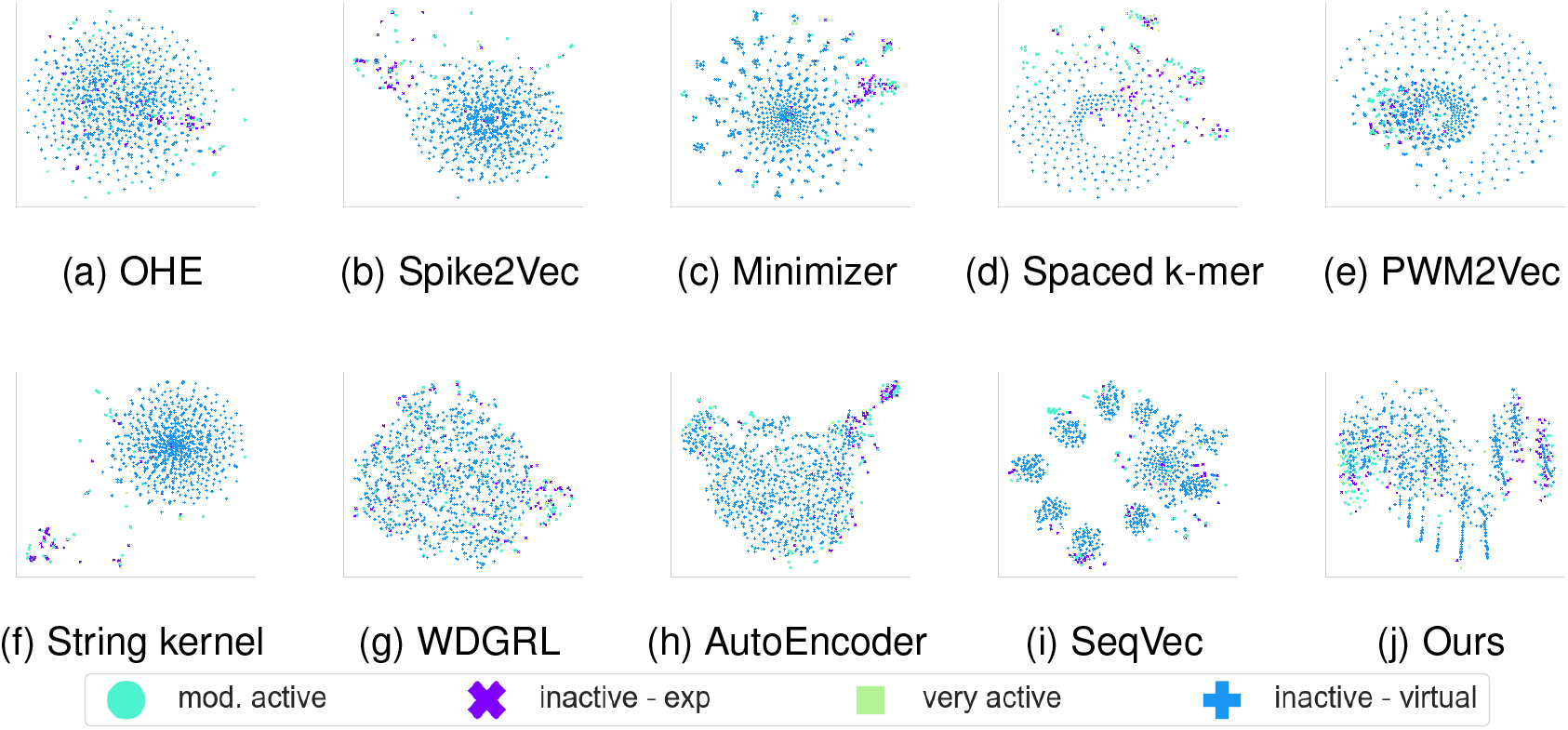
t-SNE plots for (**Breast Cancer Data**) for different structure embeddings. The figure is best seen in color.

### 4.3 Baseline Models Detail

We selected the baseline and state-of-the-art (SOTA) methods that represent different categories of sequence classification. These categories include feature engineering, kernel methods, neural networks, and pre-trained large language models. The feature engineering approache include One-Hot encoding (OHE) [28], Spike2Vec [4], Minimizers [20], Spaced *k*-mer [35], and PWM2Vec [5]. The kernel method includes String kernel [6] and Sinkhorn-Knopp Algorithm [3]. The neural network methods include WDGRL [34] and AutoEncoder [40]. Finally, the pre-trained large language models (LLMs) include SeqVec [23], Protein Bert [8], and TAPE [33]. Each method is summarized in Table 2.

**Table 2:**
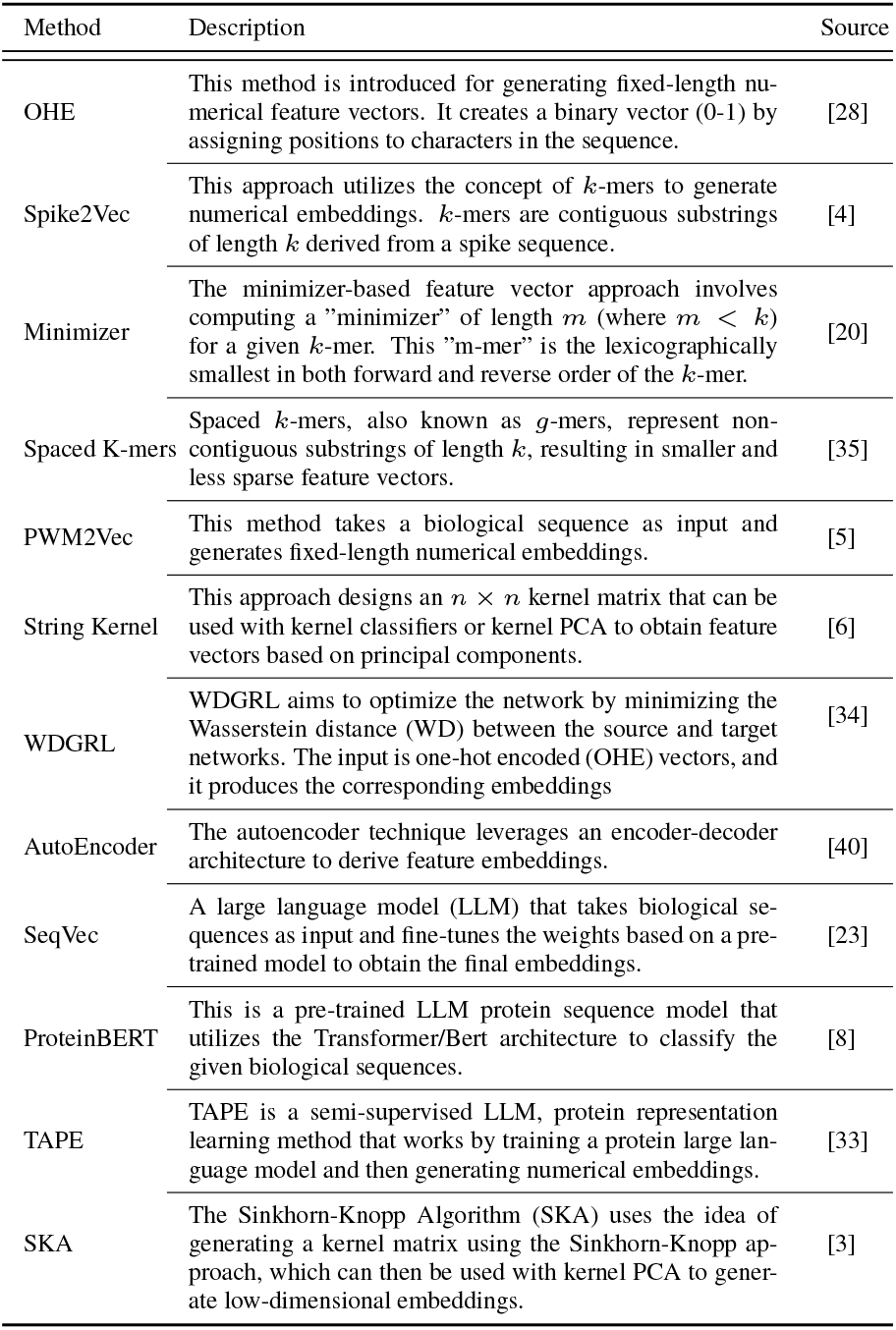
Description of different baseline models.

### 4.4 Evaluation Metrics And Classifiers

For classification tasks, we employ several classifiers including Support Vector Machine (SVM), Naive Bayes (NB), Multi-Layer Perceptron (MLP), K-Nearest Neighbors (KNN) with *K* = 5 (selected through the standard validation set approach [17]), Random Forest (RF), and Logistic Regression (LR). To assess the effectiveness, we utilize a range of evaluation metrics, including average accuracy, precision, recall, weighted metrics, and the area under the Receiver Operating Characteristic (ROC) curve (AUC).

## 5 Results And Discussion

We discuss the results of the proposed method and its comparisons with the baselines and SOTA in this section.

### 5.1 Classification Results

The classification results averaged over 5 runs for both datasets are reported in Table 3 and 4. Breast Cancer classification results show our proposed Gzip-based representation outperformed all baselines according to accuracy, precision, recall, weighted F1 score, and ROC-AUC. Even after fine-tuning the Large language models (LLM) such as SeqVec, Protein Bert, and TAPE, the proposed parameter-free method significantly outperforms the LLMs for all evaluation metrics. TAPE has better ROCAUC but it is very marginal and for other metrics our proposed method still outperforms. For classification training runtime, WDGRL with Naive Bayes performs the best due to the smaller embedding size. We observe similar results for Lung Cancer classification, where our Gzip-based method outperforms all baselines in the majority of the metrics except for ROC-AUC and F1 Macro score.

**Table 3:** Average classification results for **Breast Cancer** dataset. The best values for each metric are underlined. Overall best values are shown in bold.

| Embedding Method | ML Algo. | Acc. ↑ | Prec. ↑ | Recall ↑ | F1 (Weig.) ↑ | F1 (Macro) ↑ | ROC AUC ↑ | Training Time (sec.) ↓ |
| --- | --- | --- | --- | --- | --- | --- | --- | --- |
| OHE | SVM | 0.882 | <u>0.883</u> | 0.882 | 0.880 | 0.568 | <u>0.764</u> | 0.247 |
|  | NB | 0.844 | 0.842 | 0.844 | 0.834 | 0.451 | 0.686 | <u>0.028</u> |
|  | MLP | 0.849 | 0.868 | 0.849 | 0.857 | 0.521 | 0.742 | 7.933 |
|  | KNN | 0.609 | 0.853 | 0.609 | 0.676 | 0.395 | 0.678 | 0.069 |
|  | RF | 0.880 | 0.864 | 0.880 | 0.867 | 0.554 | 0.726 | 0.720 |
|  | LR | <u>0.888</u> | 0.880 | <u>0.888</u> | <u>0.881</u> | <u>0.575</u> | 0.755 | 0.052 |
| Spike2Vec | SVM | 0.878 | 0.870 | 0.878 | 0.868 | 0.534 | 0.727 | 8.342 |
|  | NB | 0.855 | 0.844 | 0.855 | 0.842 | 0.470 | 0.695 | 0.181 |
|  | MLP | 0.756 | 0.834 | 0.756 | 0.788 | 0.441 | 0.700 | 27.214 |
|  | KNN | 0.241 | 0.298 | 0.241 | 0.212 | 0.200 | 0.550 | <u>0.133</u> |
|  | RF | <u>0.893</u> | <u>0.885</u> | <u>0.893</u> | <u>0.883</u> | **0.606** | <u>0.752</u> | 1.812 |
|  | LR | 0.890 | 0.876 | 0.890 | 0.878 | 0.557 | 0.733 | 0.092 |
| Minimizer | SVM | 0.773 | 0.808 | 0.773 | 0.784 | 0.424 | 0.668 | 3.788 |
|  | NB | 0.733 | 0.806 | 0.733 | 0.762 | 0.355 | 0.653 | 0.315 |
|  | MLP | 0.766 | 0.822 | 0.766 | 0.788 | 0.441 | 0.689 | 28.805 |
|  | KNN | 0.577 | 0.807 | 0.577 | 0.635 | 0.332 | 0.616 | 0.149 |
|  | RF | 0.731 | 0.817 | 0.731 | 0.761 | 0.410 | 0.660 | 2.832 |
|  | LR | <u>0.843</u> | <u>0.836</u> | <u>0.843</u> | <u>0.833</u> | <u>0.483</u> | <u>0.690</u> | <u>0.109</u> |
| Spaced k-mer | SVM | 0.865 | 0.842 | 0.865 | 0.845 | 0.465 | 0.674 | 127.502 |
|  | NB | 0.469 | <u>0.865</u> | 0.469 | 0.599 | 0.359 | 0.607 | 4.302 |
|  | MLP | 0.420 | 0.851 | 0.420 | 0.497 | 0.341 | 0.624 | 204.755 |
|  | KNN | 0.276 | 0.460 | 0.276 | 0.253 | 0.216 | 0.559 | <u>1.036</u> |
|  | RF | 0.348 | 0.864 | 0.348 | 0.363 | 0.304 | 0.598 | 38.087 |
|  | LR | <u>0.869</u> | 0.848 | <u>0.869</u> | <u>0.851</u> | <u>0.480</u> | <u>0.682</u> | 1.445 |
| PWM2Vec | SVM | 0.804 | 0.834 | 0.804 | 0.811 | 0.434 | 0.687 | 132.720 |
|  | NB | 0.826 | 0.846 | 0.826 | 0.823 | 0.408 | 0.670 | 0.887 |
|  | MLP | 0.764 | 0.841 | 0.764 | 0.791 | <u>0.462</u> | <u>0.720</u> | 36.824 |
|  | KNN | 0.199 | 0.808 | 0.199 | 0.221 | 0.190 | 0.541 | <u>0.618</u> |
|  | RF | 0.754 | <u>0.857</u> | 0.754 | 0.788 | 0.460 | 0.717 | 6.479 |
|  | LR | <u>0.839</u> | 0.819 | <u>0.839</u> | <u>0.824</u> | 0.434 | 0.670 | 16.056 |
| String Kernel | SVM | 0.808 | 0.865 | 0.808 | 0.829 | 0.503 | 0.730 | 0.316 |
|  | NB | 0.874 | 0.880 | 0.874 | <u>0.873</u> | <u>0.545</u> | <u>0.746</u> | <u>0.011</u> |
|  | MLP | 0.634 | 0.789 | 0.634 | 0.684 | 0.358 | 0.648 | 5.259 |
|  | KNN | <u>0.881</u> | 0.860 | <u>0.881</u> | 0.862 | 0.494 | 0.705 | 0.030 |
|  | RF | 0.879 | <u>0.862</u> | 0.879 | 0.861 | 0.498 | 0.697 | 1.133 |
|  | LR | 0.790 | 0.843 | 0.790 | 0.812 | 0.464 | 0.707 | 0.398 |
| WDGRL | SVM | 0.804 | 0.646 | 0.804 | 0.716 | 0.223 | 0.500 | 0.026 |
|  | NB | 0.779 | 0.718 | 0.779 | 0.735 | <u>0.305</u> | 0.535 | **0.002** |
|  | MLP | 0.778 | 0.714 | 0.778 | 0.735 | 0.301 | <u>0.536</u> | 3.588 |
|  | KNN | 0.794 | 0.715 | 0.794 | 0.730 | 0.270 | 0.518 | 0.016 |
|  | RF | 0.667 | 0.689 | 0.667 | 0.672 | 0.301 | 0.532 | 0.021 |
|  | LR | <u>0.818</u> | <u>0.825</u> | <u>0.818</u> | <u>0.749</u> | 0.301 | 0.531 | 0.006 |
| Auto-Encoder | SVM | 0.816 | 0.813 | 0.816 | 0.813 | 0.443 | 0.678 | 0.109 |
|  | NB | 0.735 | 0.754 | 0.735 | 0.738 | 0.360 | 0.630 | <u>0.009</u> |
|  | MLP | 0.806 | 0.808 | 0.806 | 0.806 | 0.451 | 0.675 | 15.740 |
|  | KNN | 0.832 | 0.802 | 0.832 | 0.804 | 0.431 | 0.645 | 0.067 |
|  | RF | <u>0.846</u> | 0.805 | <u>0.846</u> | 0.817 | 0.439 | 0.648 | 2.210 |
|  | LR | 0.834 | <u>0.820</u> | 0.834 | <u>0.826</u> | <u>0.464</u> | <u>0.684</u> | 0.693 |
| SeqVec | SVM | 0.886 | <u>0.881</u> | 0.886 | <u>0.882</u> | 0.531 | 0.735 | 80.105 |
|  | NB | 0.625 | 0.757 | 0.625 | 0.665 | 0.244 | 0.607 | 4.691 |
|  | MLP | 0.848 | 0.872 | 0.848 | 0.856 | 0.516 | <u>0.743</u> | 160.886 |
|  | KNN | 0.674 | 0.819 | 0.674 | 0.725 | 0.389 | 0.651 | 22.253 |
|  | RF | <u>0.887</u> | 0.874 | <u>0.887</u> | 0.876 | <u>0.551</u> | 0.725 | 11.360 |
|  | LR | 0.769 | 0.872 | 0.769 | 0.813 | 0.465 | 0.725 | 1816.404 |
| Protein Bert<sub>+</sub> |  | 0.893 | 0.893 | 0.893 | 0.893 | 0.602 | 0.779 | 64.849 |
| TAPE | SVM | 0.890 | 0.892 | 0.890 | 0.890 | 0.579 | 0.774 | 0.042 |
|  | NB | 0.859 | 0.888 | 0.859 | 0.870 | 0.584 | **0.795** | 0.022 |
|  | MLP | 0.886 | 0.893 | 0.886 | 0.889 | <u>0.592</u> | 0.781 | 0.926 |
|  | KNN | 0.886 | 0.875 | 0.886 | 0.879 | 0.551 | 0.745 | <u>0.021</u> |
|  | RF | 0.885 | 0.863 | 0.885 | 0.871 | 0.523 | 0.724 | 3.141 |
|  | LR | <u>0.893</u> | <u>0.894</u> | <u>0.893</u> | <u>0.894</u> | <u>0.592</u> | 0.778 | 0.530 |
| SKA | SVM | 0.840 | 0.808 | 0.840 | 0.799 | 0.425 | 0.613 | 0.347 |
|  | NB | <u>0.884</u> | <u>0.881</u> | <u>0.884</u> | <u>0.873</u> | <u>0.547</u> | <u>0.752</u> | <u>0.011</u> |
|  | MLP | 0.542 | 0.681 | 0.542 | 0.588 | 0.280 | 0.545 | 11.433 |
|  | KNN | 0.236 | 0.226 | 0.236 | 0.213 | 0.199 | 0.546 | 0.080 |
|  | RF | 0.875 | 0.848 | 0.875 | 0.858 | 0.483 | 0.708 | 1.412 |
|  | LR | 0.793 | 0.629 | 0.793 | 0.702 | 0.221 | 0.500 | 0.084 |
| Ours | SVM | 0.783 | 0.614 | 0.783 | 0.688 | 0.220 | 0.500 | 0.247 |
|  | NB | 0.587 | 0.880 | 0.587 | 0.694 | 0.453 | 0.679 | <u>0.018</u> |
|  | MLP | 0.752 | 0.662 | 0.752 | 0.696 | 0.263 | 0.518 | 2.201 |
|  | KNN | 0.783 | 0.614 | 0.783 | 0.688 | 0.220 | 0.500 | 0.271 |
|  | RF | **0.915** | **0.910** | **0.915** | **0.910** | <u>0.579</u> | <u>0.784</u> | 0.728 |
|  | LR | 0.783 | 0.614 | 0.783 | 0.688 | 0.220 | 0.500 | 0.055 |

**Table 4:**
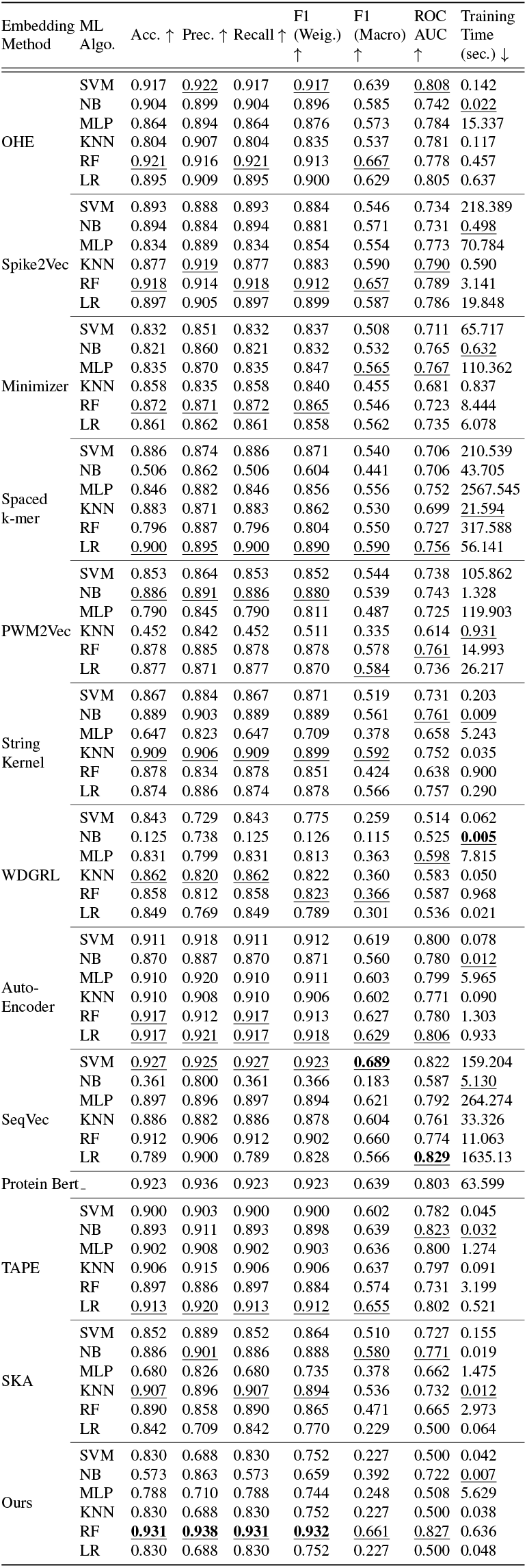
Average classification results for **Lung Cancer** dataset. The best values for each metric are underlined. Overall best values are shown in bold.

### 5.2 Class-Wise Analysis Using Heatmaps

We employ heat maps to delve deeper into the effectiveness of our proposed approach in distinguishing between various classes. These maps are created by initially calculating the pairwise cosine similarity between embeddings of different classes and then averaging the similarity values to derive a singular value for each class pair. Heatmaps are shown in Figure 4 and Figure 5 (in the appendix) for Breast Cancer and Lung Cancer data, respectively. These figures show the inter-class and intra-class similarity. The light color diagonals in all figures show a high positive intra-class similarity, depicting a strong resemblance of the ACP sequences belonging to the same category. The portions other than the diagonals in the heatmap represent inter-class similarity. The heatmaps for the proposed *k*-mer compression-based method are given in Figure 4j and Figure 5j. It can be observed that the similarity between different classes is very low, showing that our proposed embedding differentiates between classes. While the heatmaps for baselines, especially WDGRL and SeqVec, shown in Figure (4g-4i) and Figure (5g-5i) mostly have positive high similarity between all the classes showing zero to very weak differentiation between different classes of ACPs in the embedding feature space as they are very close to each other.

### 5.3 Class-Wise Analysis Using Bar Plots

We use bar plots to analyze the values in different embeddings and compute their kernel values to observe which ones have better class-wise separations. An example of a pair of embeddings (using the Spike2Vec approach) belonging to the same class and a pair of embeddings belonging to a different class is shown in Figure 6 for randomly selected pairs for the breast cancer dataset (in the appendix). In Figure 6 (a) and (b), we can observe the Spike2Vec-based *k*-mers spectrum for the “mod. active” label. As they belong to the same class, we expect the pairwise Gaussian kernel value to be as big as possible. The Gaussian kernel value for Figure 6 (a) and (b) is 0.98412732 using *k*-mers spectrum embedding, while for our proposed *k*-mer Compression-based embedding, the Gaussian kernel value is 0.99999999, hence showing that the proposed method captures the similarity among same class better. Similarly, Figure 6 (c) and (d) represent *k*-mers spectrum embeddings for different classes (selected randomly), and we can expect the Gaussian kernel value to be smaller, which we observed in the case of the proposed *k*mers compression-based approach compared to Spike2Vec-based *k*-mers spectrum.

For the Lung cancer ACP dataset, we can observe similar behavior in Figure 7 (in the appendix). The Gaussian kernel value for Figure 7 (a) and (b), which represents the label “very active”, is 0.9884221 for Spike2Vec, while for *k*-mer Compression, it is 0.9999999 (larger value is better). Similarly, the Gaussian kernel value for Figure 7 (c) and (d) is 0.9999060 for Spike2Vec, which represents the label “inactive-exp” and “very active”, while for the *k*-mer Compression, the Gaussian kernel value is 0.99989160 (a smaller kernel value is better). This behavior also shows that the embeddings generated using the proposed *k*-mers compression-based approach better show the inter-class-based separations and intra-class-based closeness.

### 5.4 Statistical Significance

We conducted a student t-test, the *p*-values were derived from the average and standard deviations obtained from five independent random runs. It is noteworthy that all computed *p*-values were below 0.05, indicating the statistical significance of the results. This observation can be attributed to the generally low standard deviation (SD) values across the data. The SD results for breast cancer and lung cancer datasets are in Table 5 and Table 6 (in the appendix), respectively. We can observe that the SD values of our method and most of the baselines are very small, which shows that the results are stable.

From the overall average and SD results, we can see that our method outperforms the SOTA for predictive performance on the real-world Anti-Cancer Peptides (ACPs) sequence datasets. The results demonstrate the effectiveness of our Gzip-based representation method over various baselines across multiple evaluation metrics. This superiority is particularly notable in terms of accuracy, precision, recall, weighted F1 score, and ROC-AUC. Such consistent outperformance indicates the robustness and reliability of our approach across different classification tasks. Despite fine-tuning large language models (LLMs) like SeqVec and Protein Bert, our parameter-free method outperforms them across all evaluation metrics. This observation is particularly intriguing given the complexity and expressiveness of LLMs, suggesting that domain-specific representations, such as the Gzip-based method proposed, can provide more tailored and effective solutions for sequence analysis. Also, it can help biologists better understand Cancer Biology and come up with improved cancer prediction and treatment methods.

While the proposed method excels in classification accuracy, computational efficiency is crucial for large-scale applications. Despite competitive performance, methods like SeqVec and Spaced *k*-mers may require more computational resources, posing practical limitations. Class-wise similarity plots reveal strong intra-class similarities and low interclass similarities, indicating effective differentiation and validating the discriminative power of the proposed embedding method. These promising results open up avenues for its application in various bioinformatics and medical domains beyond cancer classification. Such as facilitating the development of diagnostic tools, drug discovery pipelines, and personalized medicine approaches.

## 6 Conclusion

We propose a lightweight and efficient compression-based method involving *k*-mer strategy and NLP-based encoding for classifying Anti-Cancer Peptide sequences. Our method achieves SOTA performance without the need for parameter tuning or pre-trained models. The compression-based model successfully overcame the limitations of neural networkbased methods, offering improved accessibility and computational efficiency, especially in low-resource scenarios. In future research, we will explore the applications of our model in other biological domains and investigate ways to optimize the method for specific biological datasets.

## References

[1] S. Ahmed, R. Muhammod, et al., Acp-mhcnn: An accurate multi-headed deep-convolutional neural network to predict anticancer peptides, Scientific reports, 11 (2021), p. 23676.

[2] S. Akbar, M. Hayat, et al., cacp-deepgram: classification of anticancer peptides via deep neural network and skipgram-based word embedding model, Artificial intelligence in medicine, 131 (2022), p. 102349.

[3] S. Ali, T. E. Ali, T. Murad, H. Mansoor, and M. Patterson, Molecular sequence classification using efficient kernel based embedding, Information Sciences, 679 (2024), p. 121100.

[4] S. Ali, B. Bello, P. Chourasia, R. T. Punathil, P.-Y. Chen, I. U. Khan, and M. Patterson, Virus2vec: Viral sequence classification using machine learning, arXiv preprint 2304.12328, (2023).

[5] S. Ali et al., PWM2Vec: An efficient embedding approach for viral host specification from coronavirus spike sequences, Biology, 11 (2022), p. 418.

[6] S. Ali, B. Sahoo, M. A. Khan, A. Zelikovsky, I. U. Khan, and M. Patterson, Efficient approximate kernel based spike sequence classification, IEEE/ACM Transactions on Computational Biology and Bioinformatics, (2022).

[7] D. Azevedo, A. M. Rodrigues, H. Canhão, A. M. Carvalho, and A. Souto, Zgli: A pipeline for clustering by compression with application to patient stratification in spondyloarthritis, Sensors, 23 (2023).

[8] N. Brandes, D. Ofer, Y. Peleg, N. Rappoport, and M. Linial, Proteinbert: A universal deep-learning model of protein sequence and func., Bioinformatics, 38 (2022).

[9] M. Burdukiewicz et al., Cancergram: An effective classifier for differentiating anticancer from antimicrobial peptides, Pharmaceutics, 12 (2020), p. 1045.

[10] P. Charoenkwan, W. Chiangjong, et al., Improved prediction and characterization of anticancer activities of peptides using a novel flexible scoring card method, Scientific reports, 11 (2021), p. 3017.

[11] L. Chen, Z. Hu, et al., Deep2pep: A deep learning method in multi-label classification of bioactive peptide, Computational Biology and Chemistry, (2024), p. 108021.

[12] W. Chen, H. Ding, P. Feng, H. Lin, and K.-C. Chou, iacp: a sequence-based tool for identifying anticancer peptides, Oncotarget, 7 (2016), p. 16895.

[13] Z. Chen, P. Zhao, F. Li, et al., ifeature: a python package and web server for features extraction and selection from protein and peptide sequences, Bioinformatics, 34 (2018), pp. 2499–2502.

[14] W. Chiangjong, S. Chutipongtanate, and S. Hongeng, Anticancer peptide: Physicochemical property, functional aspect and trend in clinical application, International journal of oncology, 57 (2020), pp. 678–696.

[15] P. Chourasia et al., Enhancing t-sne performance for biological sequencing data through kernel selection, in ISBRA, Springer, 2023, pp. 442–452.

[16] J. E. Cronan, The chain-flipping mechanism of acp (acyl carrier protein)-dependent enzymes appears universal, Biochemical Journal, 460 (2014), pp. 157–163.

[17] P. Devijver and J. Kittler, Pattern recognition: A statistical approach, in London, GB: Prentice-Hall, 1982, pp. 1–448.

[18] Z. Du, X. Ding, Y. Xu, and Y. Li, Unidl4biopep: a universal deep learning architecture for binary classification in peptide bioactivity, Briefings in Bioinformatics, 24 (2023), p. bbad135.

[19] E. Fazal, M. S. Ibrahim, et al., Anticancer peptides classification using kernel sparse representation classifier, IEEE Access, 11 (2023), pp. 17626–17637.

[20] S. Girotto, C. Pizzi, et al., Metaprob: accurate metagenomic reads binning based on probabilistic sequence signatures, Bioinformatics, 32 (2016), pp. i567–i575.

[21] GRISONI et al., ‘de novo design of anticancer peptides by ensemble artificial neural networks’, ‘Journal of Molecular Modeling’, ‘25’ (‘2019’), p. ‘112’.

[22] Z. Hajisharifi, M. Piryaiee, et al., Predicting anticancer peptides with chou’s pseudo amino acid composition and investigating their mutagenicity via ames test, Journal of theoretical biology, 341 (2014), pp. 34–40.

[23] M. Heinzinger et al., Modeling aspects of the language of life through transfer-learning protein sequences, BMC bioinformatics, 20 (2019), pp. 1–17.

[24] K.-Y. Huang, Y.-J. Tseng, et al., Identification of subtypes of anticancer peptides based on sequential features and physicochemical properties, Scientific reports, 11 (2021), p. 13594.

[25] L. Jiang, N. Sun, Y. Zhang, X. Yu, and X. Liu, Bioactive peptide recognition based on nlp pre-train algorithm, IEEE/ACM Transactions on Computational Biology and Bioinformatics, (2023).

[26] Z. Jiang et al., Low-resource” text classification: A parameter-free classification method with compressors, in Findings of the Association for Computational Linguistics: ACL 2023, 2023, pp. 6810–6828.

[27] Z. H. Kilimci and M. Yalcin, Acp-esm: A novel framework for classification of anticancer peptides using protein-oriented transformer approach, arXiv preprint 2401.02124, (2024).

[28] K. Kuzmin, A. E. Adeniyi, et al., Machine learning methods accurately predict host specificity of coronaviruses based on spike sequences alone, Biochemical and Biophysical Research Communications, 533 (2020), pp. 553–558.

[29] C. Leslie, E. Eskin, et al., Mismatch string kernels for svm protein classification, Advances in neural information processing systems, (2003), pp. 1441–1448.

[30] Z. Lv, F. Cui, et al., Anticancer peptides prediction with deep representation learning features, Briefings in bioinformatics, 22 (2021), p. bbab008.

[31] S. Mantaci, A. Restivo, and M. Sciortino, Distance measures for biological sequences: Some recent approaches, Journal of Approximate Reasoning, 47 (2008), pp. 109–124.

[32] R. Nussinov, H. Jang, et al., Precision medicine and driver mutations: computational methods, functional assays and conformational principles for interpreting cancer drivers, PLoS computational biology, 15 (2019), p. e1006658.

[33] R. Rao, N. Bhattacharya, N. Thomas, Y. Duan, P. Chen, J. Canny, P. Abbeel, and Y. Song, Evaluating protein transfer learning with tape, Advances in neural information processing systems, 32 (2019).

[34] J. Shen, Y. Qu, et al., Wasserstein distance guided representation learning for domain adaptation, in AAAI conference on artificial intelligence, 2018.

[35] R. Singh, A. Sekhon, et al., Gakco: a fast gapped k-mer string kernel using counting, in Joint ECML and KDD, 2017, pp. 356–373.

[36] H. Sung, J. Ferlay, et al., Global cancer statistics 2020: Globocan estimates of incidence and mortality worldwide for 36 cancers in 185 countries, CA: a cancer journal for clinicians, 71 (2021), pp. 209–249.

[37] H. Tao, S. Shan, H. Fu, C. Zhu, and B. Liu, An augmented sample selection framework for prediction of anticancer peptides, Molecules, 28 (2023), p. 6680.

[38] L. Van Der Maaten and G. Hinton, Visualizing data using t-sne., Journal of machine learning research, 9 (2008).

[39] J.-P. Vert, H. Saigo, and T. Akutsu, Local alignment kernels for biological sequences, Kernel methods in computational biology, (2004), pp. 131–154.

[40] J. Xie, R. Girshick, and A. Farhadi, Unsupervised deep embedding for clustering analysis, in International conference on machine learning, 2016, pp. 478–487.

[41] M. Xie, D. Liu, and Y. Yang, Anti-cancer peptides: Classification, mechanism of action, reconstruction and modification, Open biology, 10 (2020), p. 200004.

[42] H.-C. Yi, Z.-H. You, et al., Acp-dl: a deep learning long short-term memory model to predict anticancer peptides using high-efficiency feature representation, Molecular Therapy-Nucleic Acids, 17 (2019), pp. 1–9.

[43] L. Yu, R. Jing, et al., Deepacp: a novel computational approach for accurate identification of anticancer peptides by deep learning algorithm, Molecular Therapy-Nucleic Acids, 22 (2020), pp. 862–870.

[44] C. Zhou, D. Peng, et al., Acp ms: prediction of anticancer peptides based on feature extraction, Briefings in Bioinformatics, 23 (2022), p. bbac462.

