## Supplementary material for "Compression and *k*-mer based Approach For Anticancer Peptide Analysis": Suppl

### Supplementary Material/Appendix

| Embed. Method | ML Algo. | Acc. | Prec. | Recall | F1 weigh. | F1 Macro | ROC-AUC | Train. runtime (sec.) |
|---|---|---|---|---|---|---|---|---|
| OHE | SVM | 0.011531 | 0.011787 | 0.011531 | 0.01115 | 0.054918 | 0.027033 | 0.107195 |
| | NB | 0.007276 | 0.020979 | 0.007276 | 0.010636 | 0.032285 | 0.028136 | 0.012552 |
| | MLP | 0.013814 | 0.020794 | 0.013814 | 0.015768 | 0.070364 | 0.043499 | 2.575324 |
| | KNN | 0.096558 | 0.03061 | 0.096558 | 0.076064 | 0.037224 | 0.030542 | 0.019224 |
| | RF | 0.012987 | 0.019807 | 0.012987 | 0.016738 | 0.051853 | 0.030463 | 0.272568 |
| | LR | 0.004575 | 0.006959 | 0.004575 | 0.006707 | 0.048569 | 0.024639 | 0.021389 |
| Spike2Vec | SVM | 0.010349 | 0.011348 | 0.010349 | 0.013018 | 0.041622 | 0.014485 | 3.667511 |
| | NB | 0.010928 | 0.013437 | 0.010928 | 0.012582 | 0.027492 | 0.017298 | 0.067697 |
| | MLP | 0.028136 | 0.024087 | 0.028136 | 0.024798 | 0.040874 | 0.028577 | 11.51348 |
| | KNN | 0.332312 | 0.35309 | 0.332312 | 0.3504 | 0.139923 | 0.096229 | 0.060751 |
| | RF | 0.021429 | 0.026304 | 0.021429 | 0.023503 | 0.048438 | 0.024178 | 0.949801 |
| | LR | 0.023354 | 0.024003 | 0.023354 | 0.02426 | 0.031341 | 0.021011 | 0.050051 |
| Minimizer | SVM | 0.040738 | 0.021255 | 0.040738 | 0.027105 | 0.05428 | 0.029876 | 1.511685 |
| | NB | 0.011899 | 0.01895 | 0.011899 | 0.011975 | 0.02604 | 0.017245 | 0.095413 |
| | MLP | 0.04162 | 0.012692 | 0.04162 | 0.029645 | 0.048739 | 0.025533 | 9.389144 |
| | KNN | 0.165575 | 0.039188 | 0.165575 | 0.120922 | 0.057088 | 0.033037 | 0.03619 |
| | RF | 0.080953 | 0.019017 | 0.080953 | 0.050143 | 0.044922 | 0.015386 | 0.554198 |
| | LR | 0.022522 | 0.024698 | 0.022522 | 0.021395 | 0.060888 | 0.033921 | 0.056083 |
| Spaced k-mer | SVM | 0.022136 | 0.026509 | 0.022136 | 0.0277 | 0.052306 | 0.036604 | 22.96033 |
| | NB | 0.019122 | 0.011564 | 0.019122 | 0.011426 | 0.033027 | 0.026134 | 1.461223 |
| | MLP | 0.179932 | 0.027147 | 0.179932 | 0.163132 | 0.072292 | 0.046386 | 19.1192 |
| | KNN | 0.340483 | 0.371043 | 0.340483 | 0.348999 | 0.167409 | 0.078905 | 0.026639 |
| | RF | 0.320428 | 0.027512 | 0.320428 | 0.328593 | 0.151037 | 0.081197 | 0.432502 |
| | LR | 0.01796 | 0.023061 | 0.01796 | 0.022747 | 0.044403 | 0.026633 | 0.044289 |
| PWM2Vec | SVM | 0.013361 | 0.022188 | 0.013361 | 0.017417 | 0.029452 | 0.023499 | 75.433989 |
| | NB | 0.014594 | 0.028826 | 0.014594 | 0.021347 | 0.023461 | 0.01848 | 0.130473 |
| | MLP | 0.035817 | 0.011475 | 0.035817 | 0.022992 | 0.021277 | 0.014342 | 7.247411 |
| | KNN | 0.068506 | 0.047985 | 0.068506 | 0.087948 | 0.033176 | 0.03148 | 0.093833 |
| | RF | 0.058133 | 0.013847 | 0.058133 | 0.040196 | 0.024099 | 0.015261 | 0.476376 |
| | LR | 0.01627 | 0.019851 | 0.01627 | 0.014441 | 0.03953 | 0.025154 | 3.789335 |
| String Kernel | SVM | 0.022385 | 0.020642 | 0.022385 | 0.02198 | 0.05209 | 0.028663 | 0.039227 |
| | NB | 0.021053 | 0.017239 | 0.021053 | 0.016954 | 0.030973 | 0.017838 | 0.001306 |
| | MLP | 0.071763 | 0.030056 | 0.071763 | 0.061252 | 0.038537 | 0.033347 | 1.707223 |
| | KNN | 0.017544 | 0.024288 | 0.017544 | 0.020255 | 0.026526 | 0.023847 | 0.009708 |
| | RF | 0.014166 | 0.01182 | 0.014166 | 0.017523 | 0.042271 | 0.027069 | 0.104601 |
| | LR | 0.010643 | 0.017807 | 0.010643 | 0.013159 | 0.038604 | 0.02602 | 0.027604 |
| WDGRL | SVM | 0.029251 | 0.047127 | 0.029251 | 0.040552 | 0.004488 | 0.000001 | 0.003755 |
| | NB | 0.040236 | 0.049838 | 0.040236 | 0.042602 | 0.017668 | 0.008055 | 0.000631 |
| | MLP | 0.031812 | 0.042893 | 0.031812 | 0.038649 | 0.027854 | 0.016427 | 0.306877 |
| | KNN | 0.023485 | 0.032871 | 0.023485 | 0.03317 | 0.010982 | 0.005075 | 0.001456 |
| | RF | 0.026883 | 0.030358 | 0.026883 | 0.036408 | 0.018898 | 0.00625 | 0.03639 |
| | LR | 0.027337 | 0.024129 | 0.027337 | 0.036731 | 0.010518 | 0.004581 | 0.000499 |
| Auto-Encoder | SVM | 0.020248 | 0.023967 | 0.020248 | 0.01991 | 0.022785 | 0.014621 | 0.011197 |
| | NB | 0.012796 | 0.031659 | 0.012796 | 0.018922 | 0.010107 | 0.014418 | 0.00063 |
| | MLP | 0.013947 | 0.017981 | 0.013947 | 0.014261 | 0.025607 | 0.012485 | 4.731397 |
| | KNN | 0.01137 | 0.013513 | 0.01137 | 0.015551 | 0.028315 | 0.018445 | 0.02596 |
| | RF | 0.010048 | 0.012584 | 0.010048 | 0.014677 | 0.016225 | 0.013071 | 0.100688 |
| | LR | 0.023062 | 0.028148 | 0.023062 | 0.024301 | 0.034111 | 0.023864 | 0.070495 |
| SeqVec | SVM | 0.009799 | 0.014657 | 0.009799 | 0.011993 | 0.059526 | 0.025807 | 9.309263 |
| | NB | 0.101208 | 0.061534 | 0.101208 | 0.047636 | 0.030714 | 0.035787 | 0.462643 |
| | MLP | 0.018633 | 0.00566 | 0.018633 | 0.013466 | 0.047617 | 0.028233 | 79.869113 |
| | KNN | 0.033416 | 0.027718 | 0.033416 | 0.026335 | 0.030362 | 0.016323 | 7.085517 |
| | RF | 0.014339 | 0.023789 | 0.014339 | 0.019627 | 0.084607 | 0.041548 | 1.086938 |
| | LR | 0.038708 | 0.027397 | 0.038708 | 0.034943 | 0.040742 | 0.020593 | 97.796897 |
| Protein Bert | - | 0.004001 | 0.006785 | 0.004001 | 0.005311 | 0.050968 | 0.018805 | 1.540009 |
| TAPE | SVM | 0.01915 | 0.01929 | 0.01915 | 0.01898 | 0.04191 | 0.02504 | 0.00983 |
| | NB | 0.00974 | 0.01302 | 0.00974 | 0.01047 | 0.05674 | 0.03472 | 0.00284 |
| | MLP | 0.01638 | 0.01495 | 0.01638 | 0.01499 | 0.03830 | 0.02766 | 0.12460 |
| | KNN | 0.01395 | 0.01299 | 0.01395 | 0.01336 | 0.03549 | 0.01997 | 0.00146 |
| | RF | 0.00628 | 0.00778 | 0.00628 | 0.00596 | 0.02313 | 0.01446 | 0.16484 |
| | LR | 0.00728 | 0.00950 | 0.00728 | 0.00604 | 0.03869 | 0.01830 | 0.02989 |
| SKA | SVM | 0.01657 | 0.02796 | 0.01657 | 0.02228 | 0.02139 | 0.01241 | 0.01643 |
| | NB | 0.02076 | 0.02373 | 0.02076 | 0.02550 | 0.05729 | 0.02065 | 0.00296 |
| | MLP | 0.10819 | 0.02261 | 0.10819 | 0.08848 | 0.04652 | 0.03254 | 1.47346 |
| | KNN | 0.32776 | 0.36370 | 0.32776 | 0.35196 | 0.14066 | 0.09836 | 0.02555 |
| | RF | 0.01850 | 0.02784 | 0.01850 | 0.02415 | 0.01508 | 0.00763 | 0.11076 |
| | LR | 0.02134 | 0.03403 | 0.02134 | 0.02947 | 0.00330 | 0.00000 | 0.02013 |
| Ours | SVM | 0.03172 | 0.04972 | 0.03172 | 0.04349 | 0.00499 | 0.00000 | 0.01692 |
| | NB | 0.02583 | 0.03007 | 0.02583 | 0.02812 | 0.04301 | 0.03319 | 0.00259 |
| | MLP | 0.03675 | 0.04114 | 0.03675 | 0.03880 | 0.02858 | 0.01260 | 0.56247 |
| | KNN | 0.03172 | 0.04972 | 0.03172 | 0.04349 | 0.00499 | 0.00000 | 0.02346 |
| | RF | 0.02097 | 0.02562 | 0.02097 | 0.02307 | 0.03544 | 0.02136 | 0.06625 |
| | LR | 0.03172 | 0.04972 | 0.03172 | 0.04349 | 0.00499 | 0.00000 | 0.00660 |

Table 5: Standard Deviation results for different models and algorithms for **Breast Cancer dataset**.

| Embed. Method | ML Algo. | Acc. | Prec. | Recall | F1 weigh. | F1 Macro | ROC-AUC | Train. runtime (sec.) |
|---|---|---|---|---|---|---|---|---|
| OHE | SVM | 0.002021 | 0.007971 | 0.002021 | 0.006482 | 0.039567 | 0.019805 | 0.04823 |
| | NB | 0.015214 | 0.024406 | 0.015214 | 0.021262 | 0.075804 | 0.043765 | 0.005193 |
| | MLP | 0.020446 | 0.016696 | 0.020446 | 0.018163 | 0.029846 | 0.019008 | 5.026278 |
| | KNN | 0.043661 | 0.016569 | 0.043661 | 0.028037 | 0.074342 | 0.028918 | 0.112521 |
| | RF | 0.020144 | 0.023035 | 0.020144 | 0.023632 | 0.075717 | 0.039408 | 0.087319 |
| | LR | 0.014434 | 0.013363 | 0.014434 | 0.01251 | 0.017049 | 0.016938 | 0.08448 |
| Spike2Vec | SVM | 0.012238 | 0.010408 | 0.012238 | 0.014209 | 0.036488 | 0.012129 | 44.343796 |
| | NB | 0.010631 | 0.008584 | 0.010631 | 0.013059 | 0.028304 | 0.020643 | 0.064434 |
| | MLP | 0.06497 | 0.015847 | 0.06497 | 0.044832 | 0.087087 | 0.052361 | 17.04986 |
| | KNN | 0.009622 | 0.014845 | 0.009622 | 0.008194 | 0.037584 | 0.01774 | 0.078644 |
| | RF | 0.009901 | 0.010718 | 0.009901 | 0.012357 | 0.040264 | 0.020618 | 0.407811 |
| | LR | 0.014621 | 0.020243 | 0.014621 | 0.016492 | 0.078275 | 0.053328 | 6.213785 |
| Minimizer | SVM | 0.012676 | 0.029414 | 0.012676 | 0.022768 | 0.023177 | 0.026139 | 20.338116 |
| | NB | 0.020312 | 0.018807 | 0.020312 | 0.020425 | 0.052309 | 0.022274 | 0.159142 |
| | MLP | 0.025485 | 0.034075 | 0.025485 | 0.022113 | 0.064331 | 0.042147 | 24.388504 |
| | KNN | 0.027688 | 0.045781 | 0.027688 | 0.035391 | 0.054313 | 0.038299 | 0.303424 |
| | RF | 0.01508 | 0.016571 | 0.01508 | 0.018452 | 0.025151 | 0.014603 | 4.854459 |
| | LR | 0.01928 | 0.023834 | 0.01928 | 0.022739 | 0.051185 | 0.033718 | 3.556247 |
| Spaced k-mer | SVM | 0.016585 | 0.020636 | 0.016585 | 0.020103 | 0.042259 | 0.024659 | 92.396341 |
| | NB | 0.031027 | 0.013472 | 0.031027 | 0.030243 | 0.023515 | 0.039779 | 6.405099 |
| | MLP | 0.037029 | 0.018984 | 0.037029 | 0.024521 | 0.057553 | 0.035657 | 911.399934 |
| | KNN | 0.022657 | 0.027395 | 0.022657 | 0.030217 | 0.054174 | 0.035564 | 7.183501 |
| | RF | 0.218253 | 0.016967 | 0.218253 | 0.171836 | 0.097537 | 0.044502 | 135.81018 |
| | LR | 0.013807 | 0.014127 | 0.013807 | 0.01643 | 0.055007 | 0.028169 | 8.173818 |
| PWM2Vec | SVM | 0.013905 | 0.022905 | 0.013905 | 0.017543 | 0.021991 | 0.031867 | 57.073685 |
| | NB | 0.020446 | 0.023011 | 0.020446 | 0.020328 | 0.104405 | 0.057791 | 0.427259 |
| | MLP | 0.037631 | 0.023981 | 0.037631 | 0.0269 | 0.037485 | 0.039153 | 10.862324 |
| | KNN | 0.269229 | 0.049904 | 0.269229 | 0.234733 | 0.092367 | 0.026867 | 0.129714 |
| | RF | 0.026481 | 0.023591 | 0.026481 | 0.019819 | 0.059224 | 0.057913 | 3.830312 |
| | LR | 0.010884 | 0.016044 | 0.010884 | 0.012526 | 0.025778 | 0.027001 | 3.21367 |
| String Kernel | SVM | 0.023771 | 0.021899 | 0.023771 | 0.02153 | 0.051471 | 0.028117 | 0.015709 |
| | NB | 0.009763 | 0.006735 | 0.009763 | 0.01025 | 0.040377 | 0.021839 | 0.0016 |
| | MLP | 0.059534 | 0.023637 | 0.059534 | 0.039817 | 0.038099 | 0.028761 | 1.035114 |
| | KNN | 0.019103 | 0.016853 | 0.019103 | 0.024835 | 0.058972 | 0.039131 | 0.003138 |
| | RF | 0.021516 | 0.03948 | 0.021516 | 0.030891 | 0.053883 | 0.03688 | 0.040703 |
| | LR | 0.022807 | 0.021679 | 0.022807 | 0.020637 | 0.026275 | 0.024033 | 0.020584 |
| WDGRL | SVM | 0.009622 | 0.024224 | 0.009622 | 0.015349 | 0.033016 | 0.014778 | 0.02785 |
| | NB | 0.121463 | 0.18556 | 0.121463 | 0.165471 | 0.089582 | 0.04302 | 0.004981 |
| | MLP | 0.010884 | 0.013506 | 0.010884 | 0.010673 | 0.029377 | 0.01354 | 1.052228 |
| | KNN | 0.004207 | 0.024683 | 0.004207 | 0.007249 | 0.031635 | 0.018083 | 0.032889 |
| | RF | 0.008084 | 0.024708 | 0.008084 | 0.012541 | 0.040672 | 0.016141 | 0.610966 |
| | LR | 0.011957 | 0.047911 | 0.011957 | 0.017395 | 0.047592 | 0.021039 | 0.013061 |
| Auto-Encoder | SVM | 0.013658 | 0.018207 | 0.013658 | 0.014882 | 0.028225 | 0.024744 | 0.007403 |
| | NB | 0.018707 | 0.008592 | 0.018707 | 0.017822 | 0.063253 | 0.033631 | 0.001291 |
| | MLP | 0.010946 | 0.012467 | 0.010946 | 0.010311 | 0.038383 | 0.025802 | 2.196278 |
| | KNN | 0.014898 | 0.018263 | 0.014898 | 0.015096 | 0.0576 | 0.018675 | 0.071613 |
| | RF | 0.007652 | 0.009304 | 0.007652 | 0.007913 | 0.043789 | 0.025377 | 0.055156 |
| | LR | 0.013457 | 0.013796 | 0.013457 | 0.013126 | 0.056485 | 0.032336 | 0.132274 |
| SeqVec | SVM | 0.007098 | 0.008111 | 0.007098 | 0.007787 | 0.038601 | 0.019308 | 35.953073 |
| | NB | 0.302329 | 0.060397 | 0.302329 | 0.308378 | 0.077244 | 0.04259 | 0.441012 |
| | MLP | 0.024671 | 0.048065 | 0.024671 | 0.035259 | 0.066469 | 0.045369 | 218.60108 |
| | KNN | 0.006063 | 0.00704 | 0.006063 | 0.008519 | 0.051643 | 0.019611 | 2.474111 |
| | RF | 0.014621 | 0.019922 | 0.014621 | 0.015998 | 0.073811 | 0.032133 | 2.410925 |
| | LR | 0.026788 | 0.013436 | 0.026788 | 0.023216 | 0.036481 | 0.032054 | 371.36839 |
| Protein Bert | - | 0.010239 | 0.015025 | 0.010239 | 0.011147 | 0.048270 | 0.025130 | 2.886544 |
| TAPE | SVM | 0.01076 | 0.01879 | 0.01076 | 0.01376 | 0.05759 | 0.03025 | 0.00528 |
| | NB | 0.03606 | 0.02679 | 0.03606 | 0.03217 | 0.07376 | 0.04583 | 0.00597 |
| | MLP | 0.01443 | 0.01535 | 0.01443 | 0.01610 | 0.06985 | 0.03449 | 0.43111 |
| | KNN | 0.00765 | 0.01248 | 0.00765 | 0.01098 | 0.05061 | 0.03160 | 0.15033 |
| | RF | 0.03269 | 0.03090 | 0.03269 | 0.03735 | 0.06646 | 0.03248 | 0.40764 |
| | LR | 0.02177 | 0.02570 | 0.02177 | 0.02219 | 0.04776 | 0.02496 | 0.03149 |
| SKA | SVM | 0.01341 | 0.02615 | 0.01341 | 0.00880 | 0.06119 | 0.02158 | 0.02917 |
| | NB | 0.00904 | 0.01156 | 0.00904 | 0.00595 | 0.05306 | 0.02391 | 0.00560 |
| | MLP | 0.03602 | 0.02117 | 0.03602 | 0.02985 | 0.04644 | 0.02656 | 0.20291 |
| | KNN | 0.01235 | 0.02528 | 0.01235 | 0.01476 | 0.03689 | 0.01828 | 0.00054 |
| | RF | 0.01679 | 0.02085 | 0.01679 | 0.02059 | 0.00938 | 0.01768 | 0.14847 |
| | LR | 0.01994 | 0.03362 | 0.01994 | 0.02814 | 0.00294 | 0.00000 | 0.00517 |
| Ours | SVM | 0.01508 | 0.02503 | 0.01508 | 0.02115 | 0.00225 | 0.00000 | 0.00252 |
| | NB | 0.02591 | 0.01670 | 0.02591 | 0.02107 | 0.02773 | 0.03019 | 0.00315 |
| | MLP | 0.01838 | 0.01588 | 0.01838 | 0.01761 | 0.01945 | 0.01031 | 0.97232 |
| | KNN | 0.01508 | 0.02503 | 0.01508 | 0.02115 | 0.00225 | 0.00000 | 0.02482 |
| | RF | 0.00800 | 0.00469 | 0.00800 | 0.01092 | 0.03314 | 0.01579 | 0.04904 |
| | LR | 0.01508 | 0.02503 | 0.01508 | 0.02115 | 0.00225 | 0.00000 | 0.00700 |

Table 6: Standard Deviation results for different models and algorithms for **Lung Cancer dataset**.

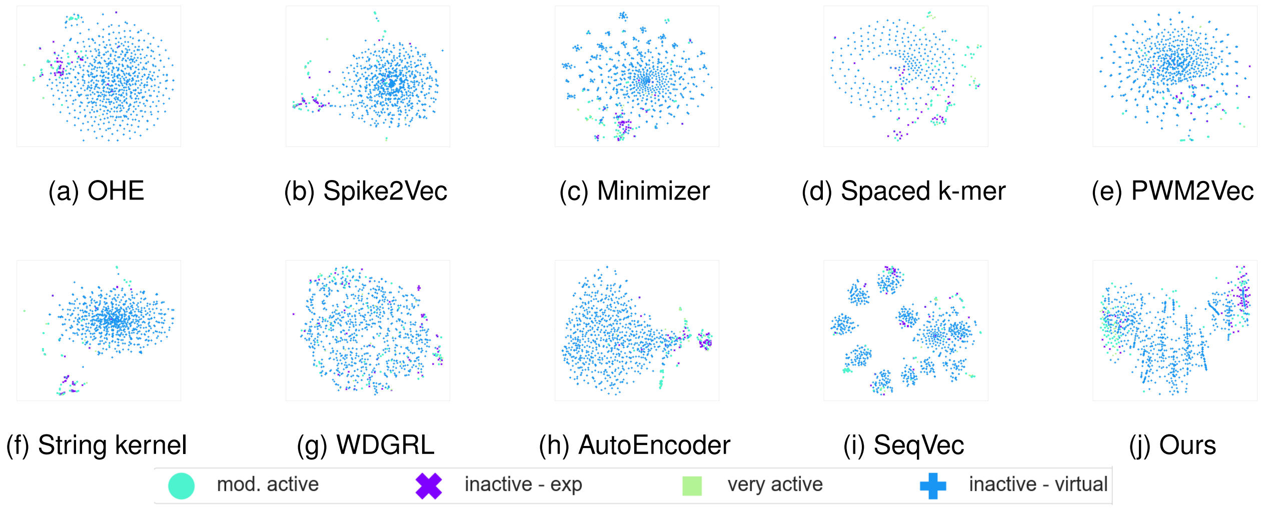


Figure 3: t-SNE plots for (**Lung Cancer Data**) for different structure embeddings. The figure is best seen in color.

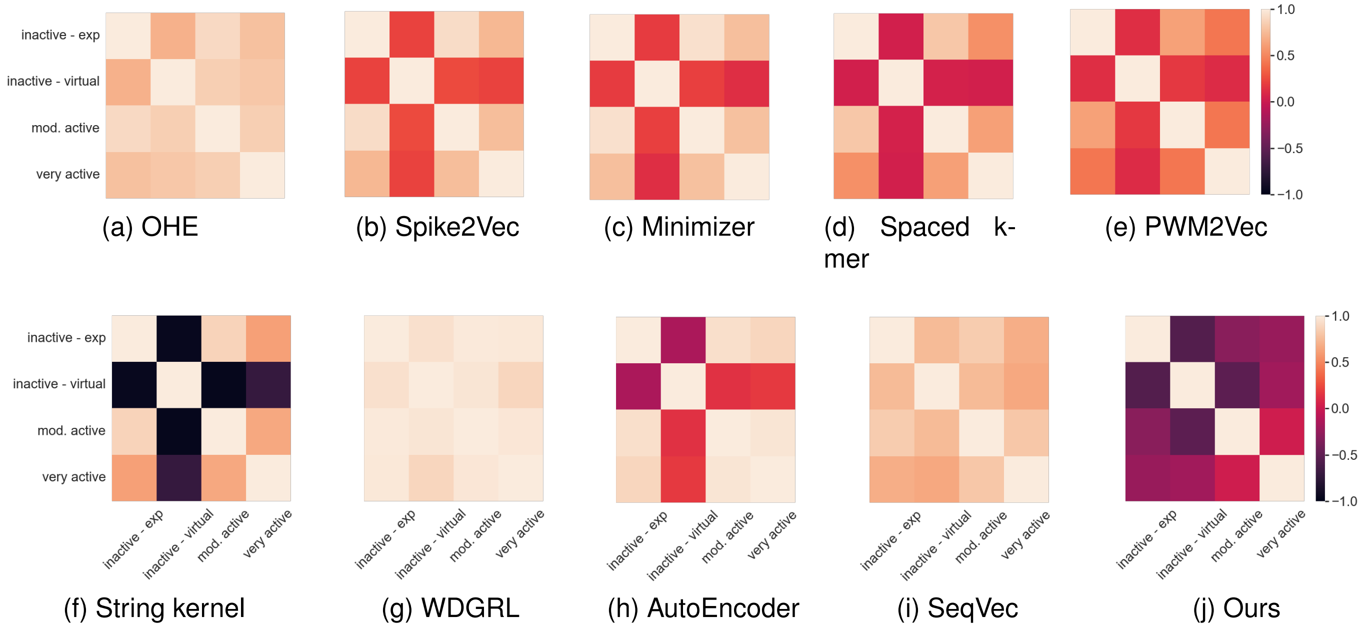


Figure 4: Heatmap plots for (**Breast Cancer Data**) for different structure embeddings. The figure is best seen in color.

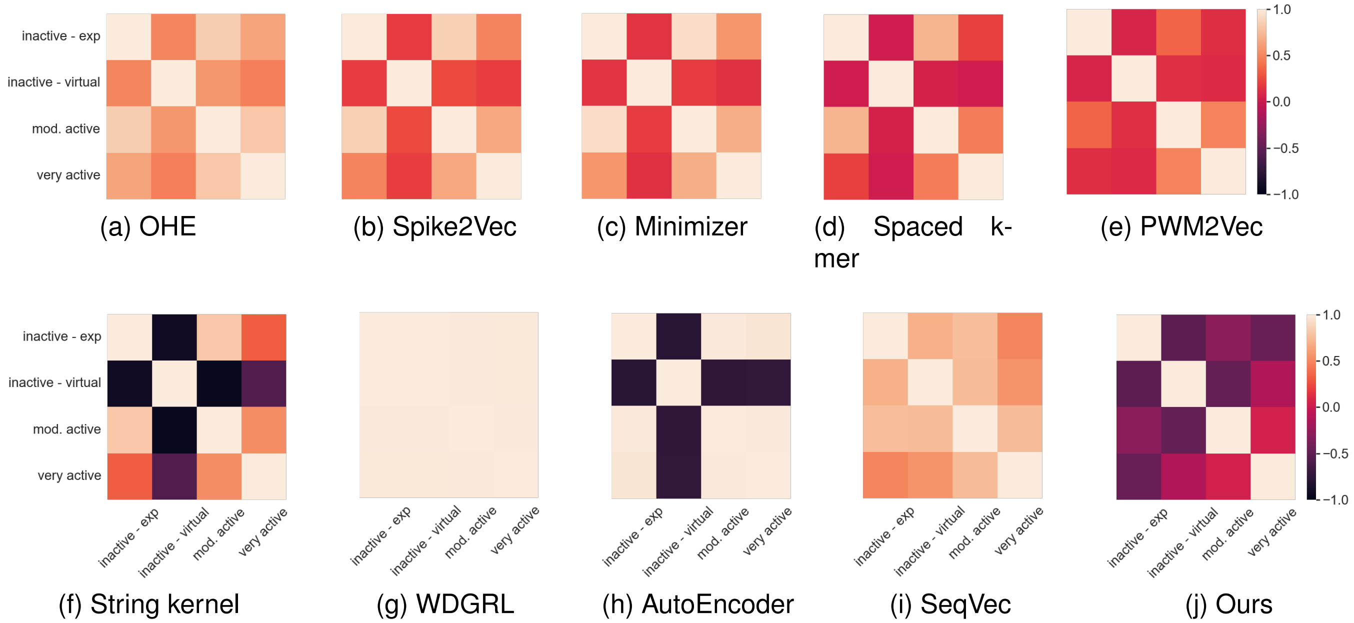


Figure 5: Heatmap plots for (**Lung Cancer Data**) for different structure embeddings. The figure is best seen in color.

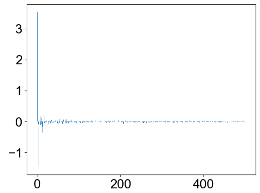

(a) very active

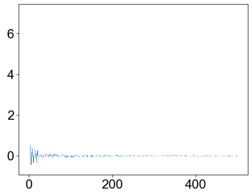

(b) very active

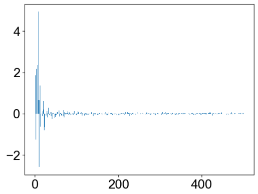

(c) inactive - exp

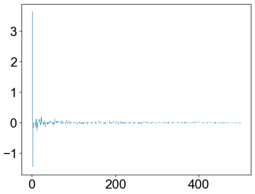

(d) very active

Figure 6: $k$-mers spectrum (i.e., Spike2Vec) embeddings of two randomly selected pairs of classes. (a) and (b) belongs to the same class, while (c) and (d) belong to different classes for the **Breast Cancer dataset**. The Gaussian kernel value for (a) and (b) is $0.98412732$ while for $k$-mer Compression is $0.99999999$ (larger value is better). The Gaussian kernel for (c) and (d) is $0.99987500$ while for the $k$-mer Compression is $0.99989129$ (smaller kernel value is better)

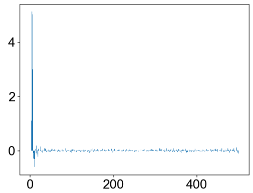

(a) very active

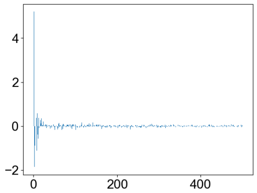

(b) very active

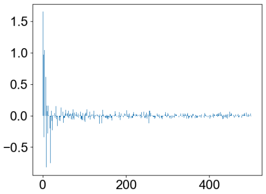

(c) inactive - exp

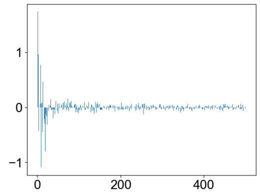

(d) very active

Figure 7: $k$-mers spectrum of two pairs of classes. (a) and (b) belongs to the same class, while (c) and (d) belong to different classes for the **Lung Cancer dataset**. The Gaussian kernel value for (a) and (b) is $0.9884221$ while for $k$-mer Compression is $0.9999999$ (larger value is better). The Gaussian kernel for (c) and (d) is $0.9999060$ while the $k$-mer Compression is $0.99989160$ (a smaller kernel value is better).
